# A Gelatin Hydrogel to Study Endometrial Angiogenesis and Trophoblast Invasion

**DOI:** 10.1101/548024

**Authors:** Samantha G. Zambuto, Kathryn B.H. Clancy, Brendan A.C. Harley

**Affiliations:** Dept. of Bioengineering University of Illinois at Urbana-Champaign Urbana, IL 61801; Dept. of Anthropology University of Illinois at Urbana-Champaign Urbana, IL 61801; Beckman Institute for Advanced Science & Technology University of Illinois at Urbana-Champaign Urbana, IL 61801; Dept. of Chemical and Biomolecular Engineering University of Illinois at Urbana-Champaign Urbana, IL 61801; Carl R. Woese Institute for Genomic Biology University of Illinois at Urbana-Champaign Urbana, IL 61801

**Keywords:** hydrogel, endometrium, trophoblast invasion, decidualization, stromal-endothelial

## Abstract

As the lining of the uterus and site of blastocyst implantation, the endometrium is a dynamic tissue that undergoes rapid cycles of growth, breakdown, and remodeling each menstrual cycle. Significant vascular remodeling is also driven by trophoblast cells that form the outer layer of the blastocyst. Trophoblast invasion and remodeling enhance blood flow to the embryo ahead of placentation. Insight into endometrial vascular remodeling and trophoblast invasion would provide key insights into endometrial physiology and cellular interactions critical for establishment of pregnancy. The objective for this study was to develop a tissue engineering platform to investigate processes of endometrial angiogenesis and trophoblast invasion in a 3D environment. We report adaptation of a methacrylamide-functionalized gelatin hydrogel that presents matrix stiffness in the range of the native tissue. Further, the hydrogel supports the formation of stable endometrial endothelial cell networks and attachment of a stratified endometrial epithelial cell layer, enables culture of a hormone-responsive stromal compartment, and provides the capacity to monitor the kinetics of trophoblast invasion. With these studies, we provide a series of techniques that will instruct researchers in the development of endometrial models of increasing complexity.

## 1. Introduction

As the lining of the uterus and site of blastocyst implantation, the endometrium is a dynamic tissue that undergoes rapid cycles of growth and breakdown over the course of the menstrual cycle. The endometrium consists of an epithelium supported by an underlying stromal layer containing endometrial glands [1]. Both the epithelium and underlying stroma undergo differentiation during the menstrual cycle to prepare for blastocyst implantation and trophoblast invasion [1]. Endometrial preparation leading up to the window of receptivity is critical for successful establishment of pregnancy. A key stage of this process is the differentiation of the endometrial stroma, or decidualization, marked by the transformation of endometrial stromal cells into specialized decidual cells that are both morphologically and functionally distinct from undifferentiated stromal cells [1, 2]. This process occurs in the presence of progesterone during the secretory phase of the menstrual cycle and continues if pregnancy occurs [1].

The endometrium is one of the only adult human tissues where non-pathological angiogenesis occurs regularly to rebuild the vascular bed [3, 4]. Further, significant vascular remodeling is driven by trophoblast cells that form the outer layer of the blastocyst in order to enhance blood flow to the embryo ahead of placentation [4–6]. During pregnancy, trophoblast cells play numerous roles, including adhesion of the blastocyst, invasion into the endometrium, anchoring the placenta, secreting hormones, modulating decidual angiogenesis, and remodeling the maternal uterine spiral arteries [6, 7]. Mutations in trophoblast function can harm maternal and fetal health. For example, insufficient endometrial vascular remodeling by trophoblast cells is believed to be a risk factor for preeclampsia [8–11]. Additionally, excessive trophoblast invasion can lead to a spectrum of placental disorders, including placenta accreta, a condition in which the placenta attaches abnormally firmly to the myometrium [7]. Despite the importance of these early events for the establishment of a successful pregnancy, initial blastocyst adhesion to the endometrial epithelium has never been observed in humans and other integral processes have been observed only from single specimens [7]. Understanding critical factors in pregnancy, including what drives trophoblast invasion and endometrial vascular remodeling, has significant value in improving our understanding of human reproduction. Therefore, strategies to monitor mechanisms associated with early trophoblast invasion and vascular remodeling in the endometrium are required to understand normal and pathological pregnancy.

Few models of the endometrium have the complexity necessary to recapitulate features of the endometrial microenvironment relevant to implantation and trophoblast invasion. These include biophysical properties, ability to monitor dynamic processes such as angiogenic events and trophoblast invasion, inclusion of a stratified structure similar to the native tissue, and selective presentation of hormonal cues. Although animal models provide the complexity of the *in vivo* environment, physiological differences in placentation between humans and animals call into question the relevance of such models for studying human pregnancy [12, 13]. While more common, traditional two-dimensional (2D) culture models do not provide the flexibility to examine endometrial cell-cell interactions in three-dimensions (3D) as they occur in the body and although 3D models could facilitate analysis of angiogenesis and trophoblast invasion, few studies to date have investigated trophoblast invasion in 3D. Further, while recent efforts have monitored trophoblast interaction with maternal vasculature [14] or the endometrial epithelium and stroma [15–17], none have developed a platform to examine the roles of the vasculature, epithelium, and stroma together. Biomaterial platforms can facilitate extended culture of cells in defined 3D microenvironments that present biophysical and biochemical properties inspired by the native tissue. Such tools are increasingly being considered to study a variety of processes relevant for endometrial physiology, including angiogenesis and invasion [18–22]. Advanced tissue engineering platforms offer the potential to enhance our understanding of core processes that influence endometrial vascular formation and trophoblast invasion through the development of increasingly complex models.

In this project, we seek to develop tissue engineering platforms to explore the relationship between biophysical, biochemical, and cellular signals from the endometrium at the maternal-fetal interface between endometrial cells and trophoblast cells. Design criteria for a such a system require integration of: (i) a hydrogel environment containing matrix stiffness relevant to the *in vivo* environment; (ii) the ability to monitor endometrial angiogenesis via culture of an endometrial perivascular niche; (iii) spatial stratified structure; (iv) capacity to vary the temporal presentation of hormonal cues (e.g. estradiol and progesterone); and (v) capability to examine trophoblast invasion in 3D (**Figure 1*a*-*b***). Here, we describe an endometrial biomaterial platform to study endometrial angiogenesis and trophoblast invasion in 3D gelatin hydrogels. The overall objective was to develop a platform to investigate endometrial angiogenesis and trophoblast invasion. Our aims were to determine how to incorporate these features into a hydrogel platform and to create a 3D trophoblast invasion assay to provide critical insight into the cellular mechanisms of endometrial vascular remodeling and trophoblast invasion.

**Figure 1.**
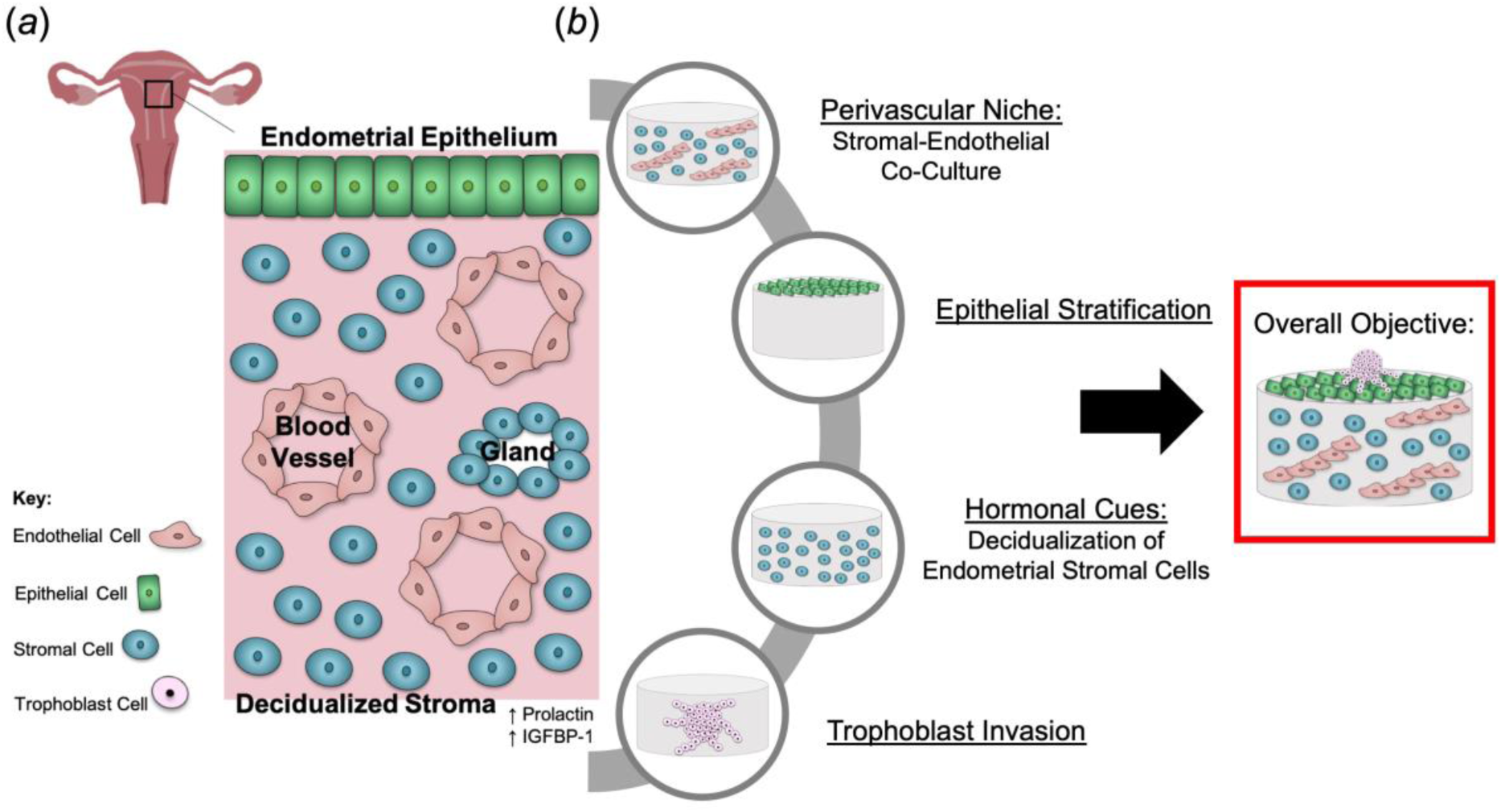
Development of a gelatin hydrogel model to recapitulate elements of the endometrial microenvironment to investigate trophoblast invasion. (*a*) *In vivo* endometrial microenvironment. (*b*) Key features to be incorporated into a gelatin hydrogel platform: endometrial perivascular niche, epithelial stratification, hormonal stimulation, and trophoblast invasion. The overall objective is to create an endometrial perivascular niche encapsulated in a gelatin hydrogel with an overlaid endometrial epithelial monolayer that can be used to study endometrial processes and trophoblast invasion.

## 2. Materials and Methods

### 2.1. Hydrogel Fabrication and Characterization

#### 2.1.1 Methacrylamide-Functionalized Gelatin (GelMA) Synthesis

GelMA was synthesized as described by Shirahama et al. [23]. Briefly, gelatin (Sigma G2500) was dissolved 10% (w/v) in carbonate buffer (Thermo Fisher 28382) at 50°C. 0.04 mL/g gelatin methacrylic anhydride (Sigma 276685) was added dropwise while stirring. The reaction was allowed to proceed for 1 hour and then was stopped by adding 40 mL/g gelatin of warm DI water. The pH of the reaction was adjusted to pH 6-7. The solution was transferred to dialysis membranes (12,000 - 14,000 molecular weight cutoff, Fisher Scientific 21-152-8), dialyzed for 1 week against DI water to remove salts and excess methacrylic acid, and subsequently lyophilized. Methacrylate degree of functionalization (DOF) was quantified via ^1^H NMR as described previously by comparing the ratio of the integrated intensity of the aromatic region (7.24 ppm) to the integrated intensity of the double bond region (5.40 and 5.64 ppm) on MestReNova (Mestrelab Research) and was determined to be approximately 66% [24, 25].

#### 2.1.2 Hydrogel Fabrication

5 wt% and 7 wt% hydrogels were prepared by dissolving lyophilized GelMA in phosphate buffered saline (PBS). After the addition of 0.1% w/v lithium acylphosphinate photoinitiator (LAP PI), prepolymer solution was cast into custom circular Teflon molds (1 mm height, 5 mm diameter) and photopolymerized for 30 s under UV light (λ=365 nm, 5.69 mW/cm^2^, AccuCure Spot System ULM-3-365).

#### 2.1.3 Compression Testing

Elastic modulus was determined by compression testing as described previously [24, 26]. Hydrogels were hydrated for 2 hours in PBS at 37°C and then subjected to unconfined compression using an Instron 5943 mechanical tester with a 5N load cell. Hydrogels were compressed with a preload of 0.5 mm/min and an extension rate of 0.1 mm/min until 30% strain was reached. Elastic modulus was determined from the slope of the linear regime (0-10% strain) of the stress-strain curve.

#### 2.3.4 Mass Swelling Ratio

5 wt% and 7 wt% hydrogels were hydrated in PBS at 37°C overnight. Swollen hydrogels were weighed, subsequently lyophilized to determine dry mass, and weighed once more. Mass swelling ratio was determined by calculating the ratio of wet polymer mass to dry polymer mass [27, 28].

### 2.2. Cell Culture

#### 2.2.1. Cell Maintenance

Human umbilical vein endothelial cells (HUVECs, Lonza C2517A, used experimentally before passage 6) were maintained in Endothelial Cell Growth Medium 2 (EGM) (PromoCell C-22211) supplemented with a SupplementPack (Promocell C-39211). Human endometrial stromal cells (HESCs, ATCC® CRL-4003, used experimentally before passage 6), normal primary human endometrial epithelial cells (EECs, LifeLine Cell Technology FC-0078, used experimentally before passage 4), and HTR-8/SVneo trophoblast cells (ATCC® CRL-3271, used experimentally before passage 6) were maintained as per the manufacturer’s instructions. HUVEC, HESC, and HTR-8/SVneo growth media were supplemented with 1% penicillin/streptomycin (Thermo Fisher 15140122). All cells were cultured in 5% CO_2_ incubators at 37°C.

#### 2.2.2 HUVEC-HESC Co-cultures

Endothelial cell networks were formed using previously described techniques by encapsulating HUVECs and HESCs in GelMA hydrogels in 2:1 and 5:1 ratios (HUVEC:HESC) with a total cell density of 3 x 10^6^ cells/mL [26, 29]. Briefly, cells were resuspended in prepolymer solution, pipetted into Teflon molds, and polymerized under UV light. Hydrogels were maintained in 48 well plates for 7 days with daily media changes. Hydrogels were maintained in phenol red-free EGM (PromoCell C-22216) supplemented with a SupplementPack and 1% penicillin/streptomycin. The fetal calf serum was replaced with charcoal-stripped fetal bovine serum (Sigma-Aldrich F6765) to reduce sex steroid hormone concentrations. The vascular endothelial growth factor (VEGF) concentration in this medium was 0.5 ng/mL.

#### 2.2.3 HESC Decidualization

Decidualization was induced by culturing HESCs in the presence of 1 μM medroxyprogesterone acetate (MPA; Sigma-Aldrich M1629) and 0.5 mM 8-Bromoadenosine 3’,5’-cyclic monophosphate (8-Br-cAMP; Sigma-Aldrich B5386) in EGM for 6 days [30–34]. For assays of decidualization in conventional 2D culture, HESCs were seeded in 6 well tissue culture plates at an initial seeding density of 36,000 cells/well. For assays of decidualization in 3D hydrogel culture, HESCs were embedded in 5 wt% GelMA hydrogels at a seeding density of 1 x 10^6^ cells/mL (approximately 20,000 cells/hydrogel). Decidualization was initiated when HESCs reached confluence for 2D culture and decidualization in hydrogels was initiated 1 day after seeding (n=3 samples for both 2D and 3D assays were imaged and analyzed). Endothelial cell growth medium (EGM) was employed as the base medium for all decidualization experiments as this medium was required for HUVEC-HESC co-culture experiments used to examine endothelial network formation (described below). Medium was replaced every 2 days and was collected for enzyme-linked immunosorbent assays (ELISAs) by centrifuging the media at 1,500 rpm for 10 minutes, aliquoting the supernatant, and storing it at −80°C until use. Human Prolactin DuoSet ELISAs (R&D DY682) and Human IGFBP-1 (insulin-like growth factor binding protein 1) DuoSet ELISAs (R&D DY871) were performed as per the manufacturer’s instructions. Samples were tested in duplicate and optical density measurements were taken at 450 nm and 540 nm using a BioTek Synergy HT Plate Reader and Gen5 software (BioTek Instruments, Inc.). Optical density was determined by subtracting the optical density at 540 nm from the optical density at 450 nm, averaging sample duplicates, and subtracting the optical density of the average of duplicate blanks (reagent diluent). OriginPro 2018b (Origin Lab) was used to fit the standard curve to a four parameter logistic curve and optical density data was inputted into the equation to extrapolate sample concentration. Cumulative release of prolactin and IGFBP-1 was calculated by adding the total concentrations from previous time points. Control data are not shown because the prolactin and IGFBP-1 concentrations in these samples fell below the detection limit of the ELISA kits.

#### 2.2.4 Epithelial Monolayer Formation

EECs were grown to confluence, passaged, and seeded at a density of 4 x 10^5^ cells/mL on 5 wt% GelMA hydrogels cast in Ibidi angiogenesis μ slides [35]. Cells were allowed to attach for 6 hours and then hydrogels were washed once with cell growth medium to remove any unattached cells. Cells were fixed 6 hours and 3 days after seeding and imaged in brightfield using a Leica DMI 4000 B microscope (Leica Microsystems).

#### 2.2.5 Trophoblast Invasion Assay

A trophoblast invasion assay was developed by embedding individual spheroids into GelMA hydrogels and observing invasion of cells into the surrounding gel matrix [36, 37], adapting methods previously described by our lab to investigate cancer cell invasion [38–40]. 2,000, 4,000, 6,000, and 8,000 HTR-8/SVneo cells were seeded into round bottom plates (Corning 4515) for 48 hours in CO_2_ incubators to create spheroids. Individual spheroids were pipetted onto prepolymer solution cast into Teflon molds. Prepolymer solution was polymerized under UV light. 8,000 cell spheroids were chosen for subsequent invasion assays due to ease of handling. Hydrogels embedded with spheroids were maintained in cell growth medium for 7 days and imaged daily. Medium was replaced on Day 3.

### 2.3 Imaging Techniques

#### 2.3.1 Immunofluorescent Staining

Immunofluorescent staining was used to visualize endothelial cell networks as described previously [26]. Briefly, hydrogels were fixed in formalin (Sigma-Aldrich HT501128) and washed 3 times with PBS. All subsequent steps were performed at room temperature on a shaker unless otherwise noted. Cells were permeabilized for 15 minutes in a 0.5% Tween 20 (Fisher Scientific BP337) solution in PBS, blocked for 1 hour in a 2% bovine serum albumin (Sigma-Aldrich A4503) and 0.1% Tween 20 solution in PBS. Hydrogels were incubated in a 1:100 dilution mouse anti-human CD31 primary antibody (Agilent IS61030-2) overnight at 4°C. Hydrogels were subsequently washed in a 0.1% Tween 20 solution in PBS for 20 minutes 4 times and incubated in a 1:500 dilution of goat anti-mouse secondary antibody (Thermo Fisher A-11001) protected from light overnight at 4°C. Hydrogels were washed in a 0.1% Tween 20 solution in PBS for 20 minutes 4 times, incubated for 30 minutes with a 1:2000 dilution of Hoechst (Thermo Fisher H3570) in PBS, and washed with the 0.1% Tween 20 solution in PBS. Samples were stored in 0.1% Tween 20 solution in PBS at 4°C until imaged.

#### 2.3.2 Analysis of Endothelial Network Complexity

Endothelial cell network images were acquired using a DMi8 Yokogawa W1 spinning disk confocal microscope outfitted with a Hamamatsu EM-CCD digital camera (Leica Microsystems). A total of 6 250 μm z-stacks with a 10 μm step size were taken for each hydrogel. As described previously [26], TubeAnalyst (IRB Barcelona), a macro available for Fiji [41], was used to quantify total vessel length, total number of junctions, total number of branches, and average branch length in 3D z-stacks. Total number of junctions and branches and total vessel length were normalized to the image volume. TubeAnalyst results were visualized using Volume Viewer and were compared manually to maximum intensity projections of each image. Regions of the skeleton that corresponded to endothelial cell sheets or image artifacts were removed.

#### 2.3.3 Analysis of Trophoblast Invasion

HTR-8/SVneo spheroids were imaged daily using a Leica DMI 4000 B microscope (Leica Microsystems). To quantify initial spheroid diameter at the day of embedding (day 0), each spheroid was manually traced using Fiji and measured using the measure tool. The perimeter values for each spheroid were measured a total of 3 times and averaged. The spheroids were assumed to be circular and radius and diameter were estimated using the equation for the perimeter of a circle. Initial spheroid area was calculated using the equation for the area of a circle. For total outgrowth area measurements, the area of each outgrowing spheroid was measured via manual tracing on Fiji [37, 42]. 3 area measurements were made for each spheroid and averaged. Fold change was calculated by comparing total spheroid outgrowth area at days 3 and 7 to the initial spheroid area at day 0.

### 2.4. Statistics

Statistical analysis was performed using OriginPro 2018b (Origin Lab). Hydrogel elastic modulus and mass swelling ratio data were analyzed using a two-sample Student’s t-test. Endothelial cell network data were analyzed using a two-sample Student’s t-test (total vessel length/mm^3^ and average branch length) or two-sample Welch’s t-test (total number of junctions and branches/mm^3^). Decidualization ELISA data and spheroid outgrowth area were analyzed via one-way ANOVA. Outgrowth area fold change was analyzed via a paired t-test. Normality was determined via the Shapiro-Wilkes test. For t-test data, equality of variance was assessed using an F-test. If the sample variances were significantly different, Welch’s correction was applied. For ANOVA data, equality of variance was determined via Levene’s test. Compression testing experiments had n=18 hydrogels per condition. Mass swelling ratio was determined for n=6 hydrogels of each condition. Analysis of endothelial cell networks was performed for n=3-6 regions imaged per hydrogel for n=3 hydrogels per experimental group. For decidualization experiments, n=3 control and n=3 treated samples were imaged and analyzed via ELISA. For spheroid invasion, n=3-6 hydrogels were measured for diameter measurements. For spheroid area outgrowth, n=6 hydrogels were imaged and measured. Outlier identification and removal was performed only on endothelial cell network data for the 2:1 HUVEC:HESC ratio using Rosner’s generalized extreme Studentized deviate test in R, assuming 3 suspected outliers and α=0.05 [43]. A total of two outliers were removed from the total number of junctions/mm^3^, one outlier was removed from total number branches/mm^3^, and three outliers were removed from total vessel length/mm^3^ (**Supplemental Figure 2**). Significance was set as *p*<0.05. Results are reported as mean ± standard deviation unless otherwise noted.

**Figure 2.**
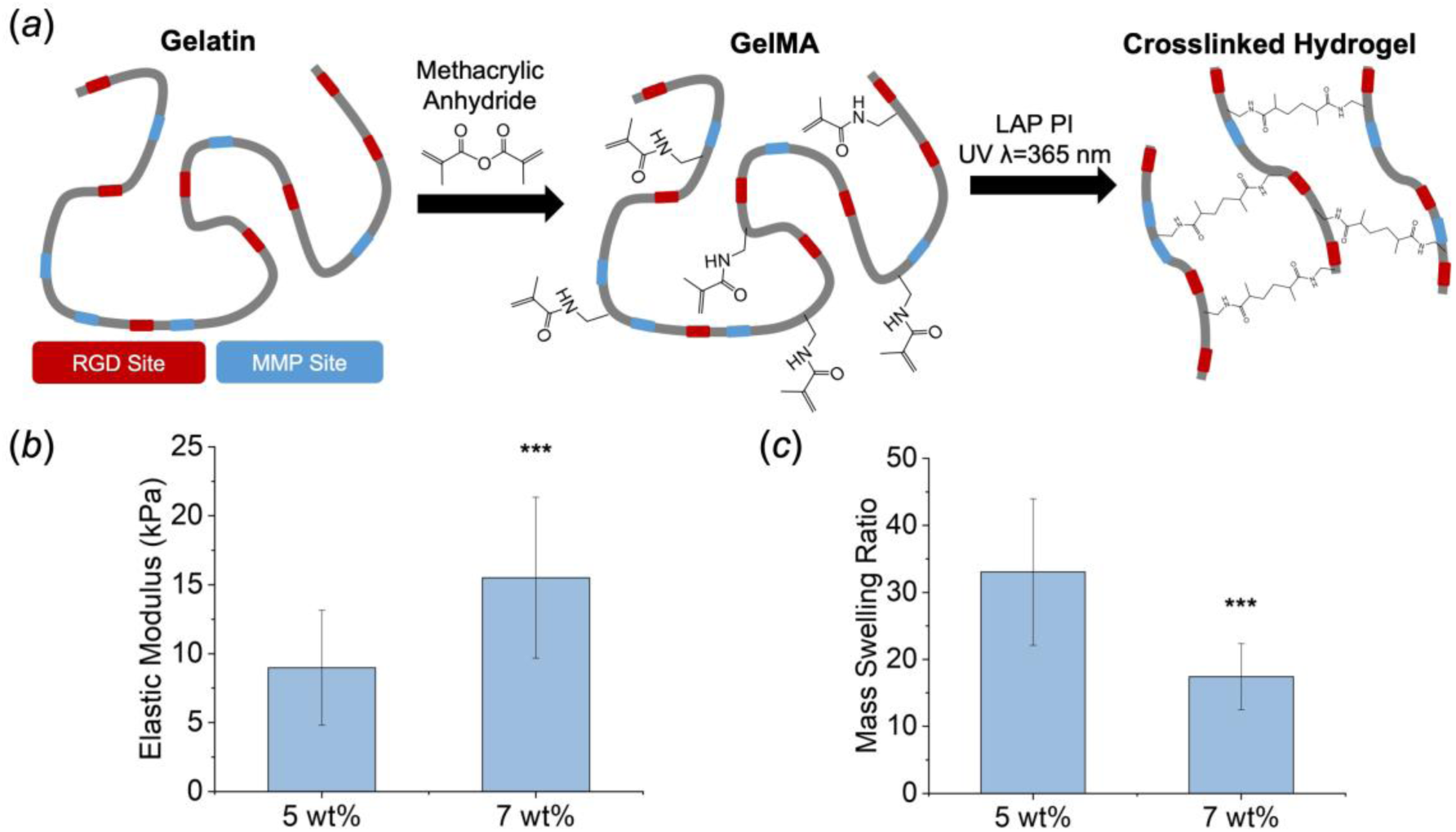
Mechanical and swelling properties of GelMA hydrogels. (*a*) Schematic of gelatin macromer, GelMA synthesis, and hydrogel formation. (*b*) Elastic moduli of 5 wt% and 7 wt% GelMA hydrogels. (*c*) Mass swelling ratio of 5 wt% and 7 wt% hydrogels. Data presented as mean ± standard deviation. ***: significant compared to 5 wt% (*p*<0.001). LAP PI: Lithium acylphosphinate photoinitiator.

## 3. Results

### 3.1. GelMA hydrogel biophysical properties and their relevance to the in vivo environment

GelMA hydrogels demonstrated a range of mechanical properties that varied with weight percent. Hydrogel elastic moduli increased significantly (*p*=4.77×10^-4^) from 5 wt% (8.97 ± 4.17 kPa) to 7 wt% (15.5 ± 5.83 kPa), respectively (**Figure 2*b***). Hydrogel mass swelling ratios were also significantly different between groups (*p*=0.0096), decreasing from 33.0 ± 10.9 to 17.4 ± 4.9 for 5 and 7 wt% variants, respectively (**Figure 2*c***). 5 wt% hydrogels were used for all subsequent cell studies as its stiffness was in a similar range as the native endometrium [44, 45].

### 3.2. Endometrial angiogenesis via culture of an endometrial perivascular niche

Co-culture of HUVECs with HESCs resulted in the formation of stable endothelial cell networks for 7 days. The endothelial cell to stromal cell ratio of cells encapsulated within the GelMA hydrogel influenced the complexity of resultant endothelial cell networks (**Figure 3*a-b***). A 2:1 HUVEC:HESC ratio led to significantly (*p*=8.79×10^-4^) increased total number of branches/mm^3^(204 ± 107) compared to a 5:1 HUVEC:HESC ratio (96 ± 42). Further, samples with a 2:1 HUVEC:HESC ratio had significantly (*p*=0.00149) increased total number of junctions/mm^3^ (65 ± 35 vs. 30 ± 15) as well as significantly (*p*=0.00167) increased total vessel length/mm^3^(12.39 ± 5.18 mm vs. 7.13 ± 3.55 mm). Average branch length was not significantly different (*p=*0.946) between the 2:1 (0.0735 ± 0.0145 mm) and 5:1 (0.0732 ± 0.0106 mm) groups.

**Figure 3.**
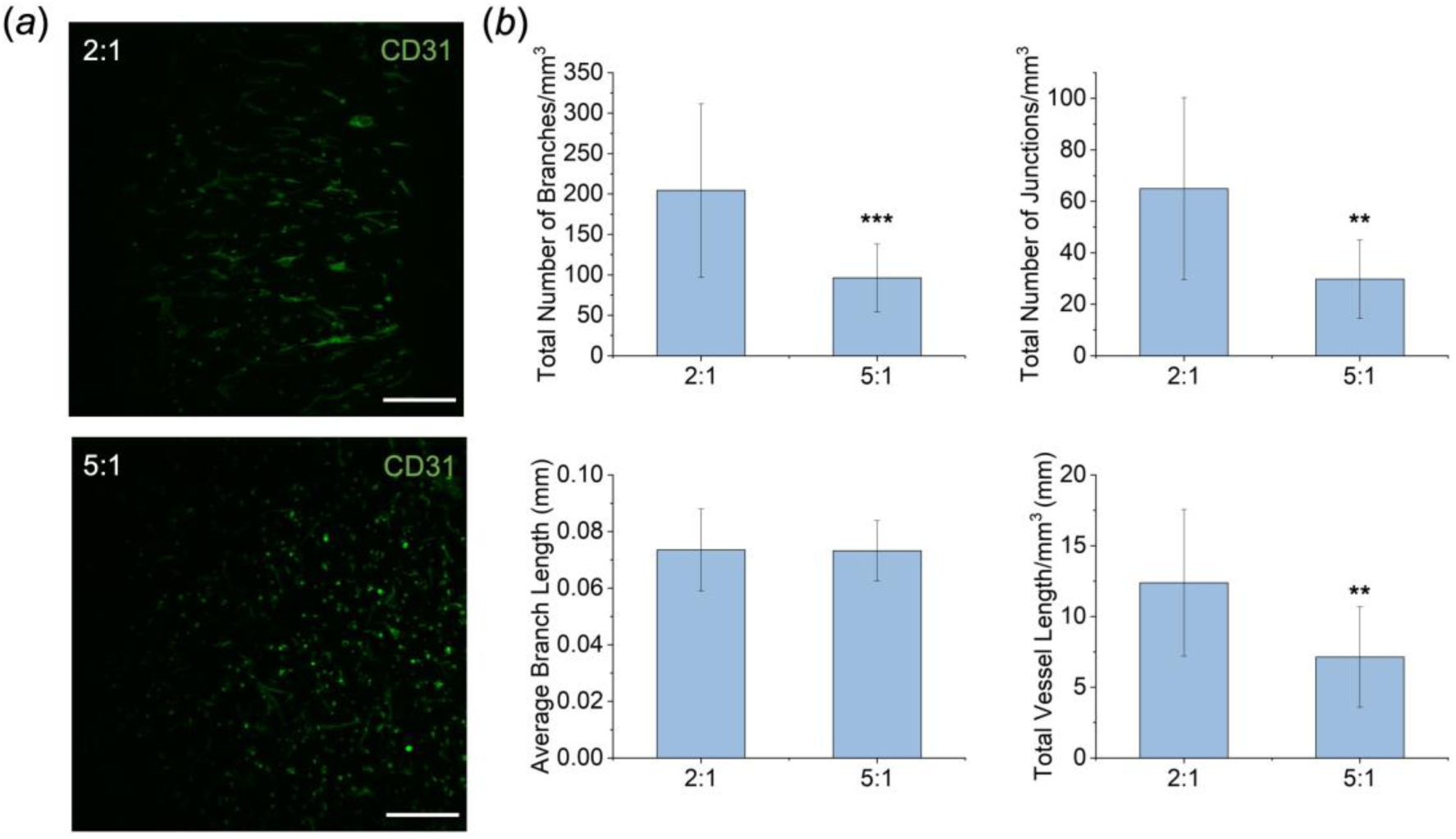
(*a*) Representative maximum intensity projection images of endothelial cell networks (green, CD31+) in hydrogels with 2:1 and 5:1 HUVEC:HESC cell seeding ratios after 7 days in culture. Images were artificially brightened for this figure to better display cells but not for analysis of network architecture. Scale bar: 250 μm. (*b*) Endothelial cell network metrics: total number of branches/mm^3^, total number of junctions/mm^3^, average branch length (mm), and total vessel length/mm^3^. These metrics were calculated from original z-stack images of 6 regions of interest per hydrogel (n=3). Data presented as mean ± standard deviation. **: significant compared to 2:1 group (*p*<0.01). ***: significant compared to 2:1 group (*p*<0.001).

### 3.3. Spatial stratification of endometrial epithelial cells

The formation of epithelial cell monolayers on top of GelMA hydrogels was explored as a function of time in culture. Primary endometrial epithelial cells adhered to GelMA hydrogels by 6 hours after seeding and remained attached to the hydrogel surface for 3 days (**Figure 4**). By 3 days in culture, epithelial cells were observed spreading along the hydrogel surface.

**Figure 4.**
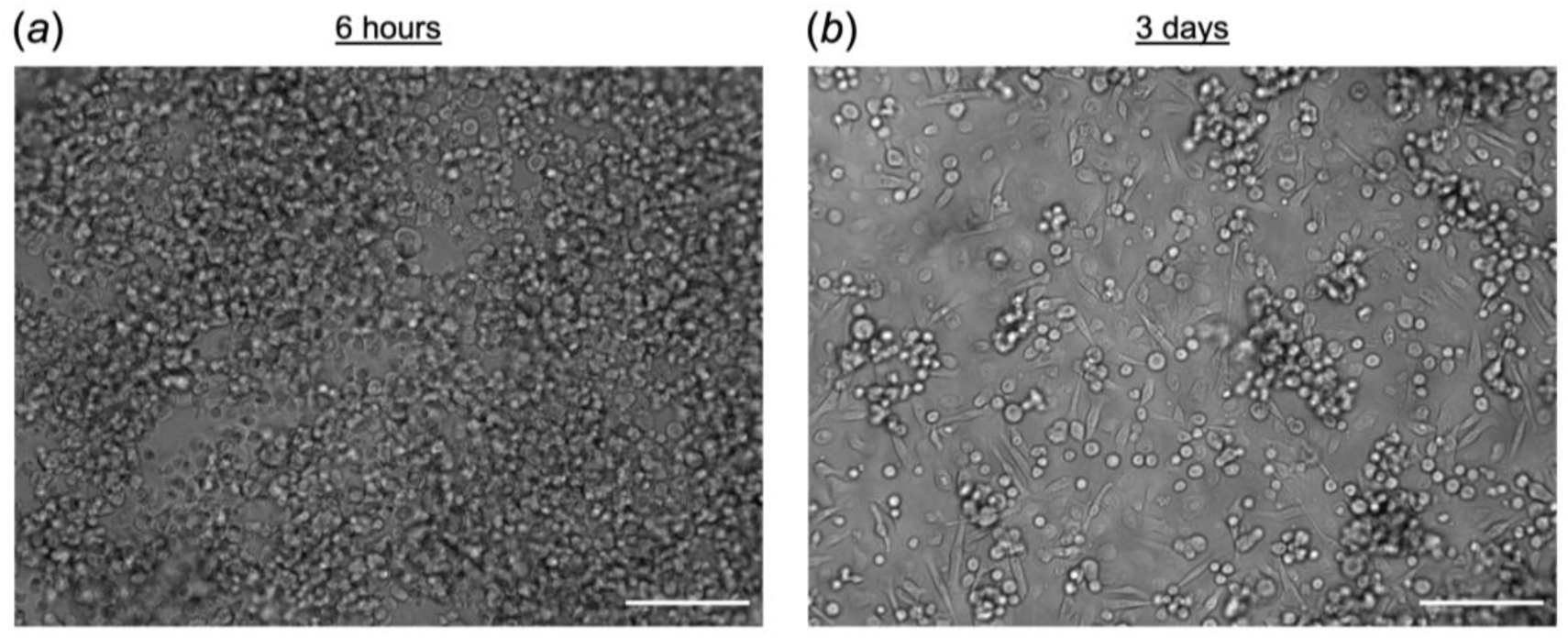
Primary endometrial epithelial cells seeded onto 5 wt% gelatin hydrogels cast in Ibidi angiogenesis μ slides. Cells were seeded onto hydrogels at an initial seeding density of 4 x 10^5^ cells/mL. (*a*) Representative bright field image of epithelial cells fixed 6 hours after seeding. (*b*) Representative bright field image of epithelial cells fixed 3 days after seeding. Scale bars: 200 μm.

### 3.4. Presentation of hormonal cues and the decidual response of endometrial stromal cells

HESC decidualization was monitored in conventional 2D cultures and 3D 5 wt% GelMA hydrogel cultures following exposure to 1 μM MPA and 0.5 mM 8-Br-cAMP. In both conventional 2D and 3D hydrogel cultures, HESCs showed characteristic epithelial-like round morphologies by 2 days that persisted for 6 days of hormonal stimulation (**Figure 5*a***). In addition, the cumulative secretion of two decidual proteins, prolactin and IGFBP-1, increased significantly (*p*<0.05) over the course of 6 days of hormonal stimulation (**Figure 5*b***) as indicated by one-way ANOVA analysis of ELISA data (Prolactin 2D: F(2,6)=389.9, *p*=4.45×10^-7^); Prolactin 3D: F(2,6)=10.3, *p*=0.011; IGFBP-1 2D: F(2,6)=88.6; *p*=3.51×10^-5^); IGFBP-1 3D: F(2,6)=75.9, *p*=5.49×10^-5^). Tukey post hoc comparisons indicated that prolactin cumulative secretion in 2D cultures significantly (*p*<0.05) differed between days 2 and 4 (*p*=4.66×10^-4^), days 2 and 6 (*p*=3.60×10^-7^), and days 4 and 6 (*p*=3.26×10^-6^). In 3D hydrogel cultures, prolactin cumulative secretion increased significantly (*p*<0.05) between days 2 and 6 (*p*=0.00963), but not between days 2 and 4 and 4 and 6. In 2D cultures, IGFBP-1 cumulative secretion significantly (*p*<0.05) differed between days 2 and 4 (*p*=0.0029), days 2 and 6 (*p*=2.80×10^-5^), and days 4 and 6 (*p*=7.02×10^-4^). In 3D hydrogel cultures, IGFBP-1 cumulative secretion significantly (*p*<0.05) differed between days 2 and 4 (*p*=7.14×10^-4^), days 2 and 6 (*p*=4.52×10^-5^), and days 4 and 6 (*p*=0.0077). Unstimulated HESCs retained their fibroblast-like morphology over the course of 6 days in culture and did not express prolactin and IGFBP-1 proteins above the detection limits of the ELISA assays used to quantify secretion. Interestingly, prolactin secretion in cell culture medium measured at each timepoint increased overtime in both 2D and 3D cultures. However, while IGFBP-1 secretion increased at each timepoint in 2D cultures, secretion levels decreased over time for 3D hydrogels cultures despite the overall increase in cumulative secretion. HESCs maintained good viability over the course of the study, as indicated by live/dead staining, and no obvious differences were observed in viability for HESCs cultured in HESC growth medium or EGM (**Supplemental Figure 1**). Taken together, these results demonstrate that decidualization of endometrial stromal cells can be induced via hormonal stimulation in 3D cultures in a manner consistent with the established 2D decidualization assays [30–34].

**Figure 5.**
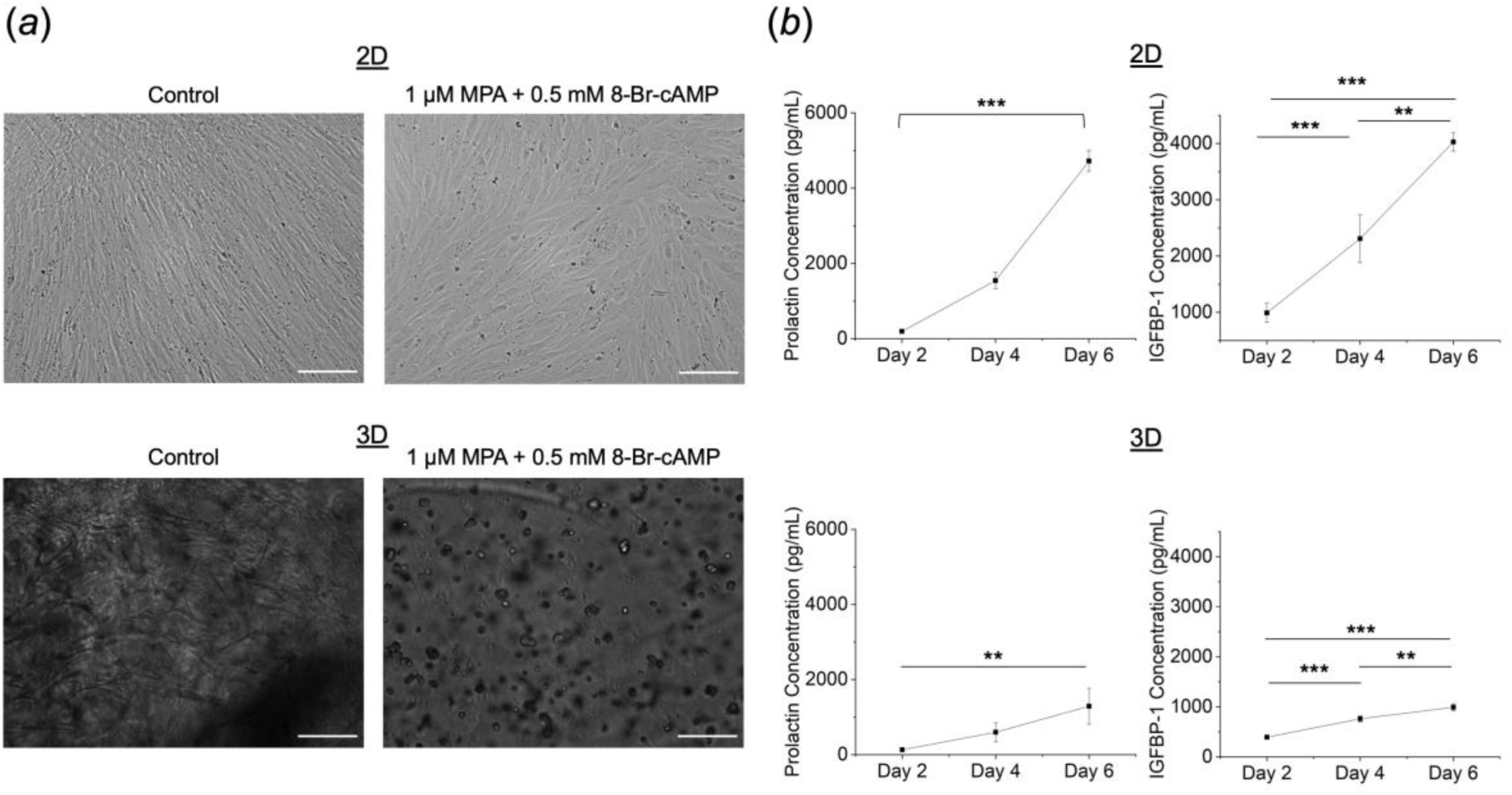
Endometrial stromal cell differentiation induced in 2D monolayers or for cells embedded in 5 wt% GelMA hydrogels via addition of 1 μM medroxyprogesterone acetate (MPA) and 0.5 mM 8-Bromo-cAMP at day 0. Medium was collected and replaced every two days for ELISA analysis. (*a*) Representative bright field images of control and treated cells in 2D or 3D hydrogel conditions at day 6 of exposure to decidualization factors. Scale bars: 200 μm. (*b*) Cumulative prolactin and IGFBP-1 concentrations for 2D and 3D cultures over 6 days. Data presented as mean ± standard deviation. **: significant (*p*<0.01); ***: significant (*p*<0.001).

### 3.5. Examination of trophoblast invasion in GelMA hydrogels

A library of HTR-8/SVneo spheroids containing 2,000, 4,000, 6,000, or 8,000 cells/spheroid displayed spheroid diameters that increased with cell density (**Table 1**). 8,000 cell spheroids were selected for invasion assays in 5 wt% GelMA hydrogels due to ease of manipulability and imaging. Trophoblast invasion into the surrounding hydrogel matrix was monitored for up to 7 days *in vitro* via serial imaging. A one-way ANOVA indicated that total outgrowth area increased significantly (*p*<0.05) over time (F(2,15)=33.7, *p*=2.81×10^-6^) (**Figure 6*b***). Tukey post hoc comparisons indicated that mean outgrowth area increased significantly (*p*=2.88×10^-6^, day 0-7; *p*=7.48×10^-5^, day 3-7) from day 0 (0.0834 ± 0.00682 mm^2^) or day 3 (0.353 ± 0.165 mm^2^) to day 7 (1.18 ± 0.385 mm^2^). No significant difference in outgrowth was observed between days 0 and Fold change in outgrowth area was subsequently calculated by comparing outgrowth areas on days 3 and 7 to initial spheroid area (Day 0, **Figure 6*c***). Fold change in outgrowth area differed significantly (*p*=6.46×10^-4^) from day 3 (3.33 ± 2.23) to day 7 (13.4 ± 5.26).

**Table 1.** Average HTR-8/SVneo spheroid diameter.

| **Spheroid Density** | **Mean Diameter ($\mu\text{m}$)** | **Standard Deviation ($\mu\text{m}$)** | **n** |
| --- | --- | --- | --- |
| 2,000 | 231.8 | 20.0 | 3 |
| 4,000 | 278.6 | 10.7 | 3 |
| 6,000 | 305.2 | 15.8 | 3 |
| 8,000 | 325.6 | 13.2 | 6 |

**Figure 6.**
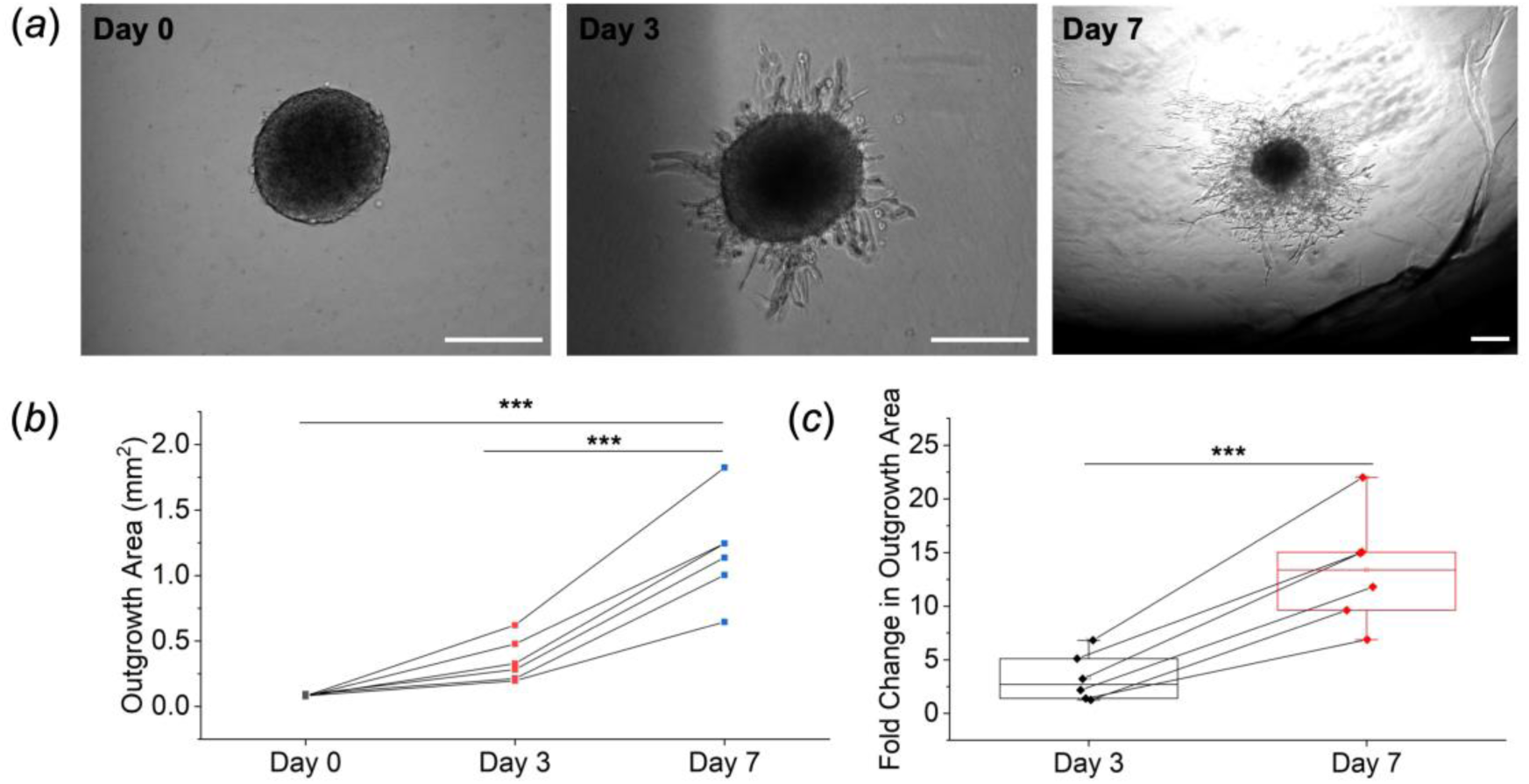
Kinetics of trophoblast invasion within a 5 wt% GelMA hydrogel. (*a*) Representative brightfield images of HTR-8/SVneo trophoblast spheroid invasion at days 0, 3, and 7. Scale bars: 250 μm. (*b*) Average outgrowth area of 8,000 HTR-8/SVneo cell spheroids at day 0, day 3, and day 7. Data presented as individual data points traced over time. (*c*) Fold change in spheroid outgrowth area at days 3 and 7. Data presented as individual data points traced over time. ***: significant (*p*<0.001).

## 4. Discussion

The objective of this study was to develop a hydrogel platform to support future mechanistic investigations of endometrial angiogenesis and trophoblast invasion. We have described a gelatin hydrogel model system that provides: (i) mechanical properties similar to native tissue; (ii) a platform to monitor endometrial angiogenesis; (iii) support of the culture of a stratified epithelial layer; (iv) capability to selectively induce endometrial stromal cell decidualization via hormonal cues; and (v) ability to perform quantitative trophoblast invasion assays. Such a platform is critical for enhancing our understanding of endometrial processes and trophoblast invasion metrics.

GelMA is an attractive material for the study of endometrial angiogenesis and trophoblast invasion for multiple reasons. Unlike synthetically-derived polymers, gelatin is denatured collagen that retains Arg-Gly-Asp (RGD) cell binding motifs and matrix metalloproteinase-sensitive degradation sites that ultimately allow for cell attachment and matrix remodeling [24, 28, 46, 47]. The addition of methacrylamide groups allows gelatin to be UV-light polymerized and hence, stable at 37°C and relatively homogeneous in terms of structure and composition [24, 28, 46, 47]. Further, the biophysical properties of GelMA can be tuned to more closely mimic the properties of native tissue. In the body, the endometrium has a stiffness ranging from 5 to 12 kPa [44, 45]. These biophysical properties informed our creation of, and down selection from, of a library of gelatin hydrogels for our model. The library we created for these studies had compressive moduli that fell within this stiffness range. We further characterized our library of hydrogels by quantifying the mass swelling ratio, an indication of the degree of crosslinking in the hydrogel [27, 47]. Mass swelling ratio decreased with hydrogel polymer weight percent, indicating that crosslinking density increased with polymer weight percent, consistent with results from other studies [27, 28, 47].

The endometrium undergoes dynamic tissue changes over the course of the menstrual cycle and during pregnancy, including vascular remodeling and angiogenesis [1, 3–5, 48]. The ability to study dynamic processes of endometrial angiogenesis would provide insight into endometrial function and vascular remodeling. Our lab [26, 29], along with other labs [49, 50], has successfully created endothelial cell networks in gelatin hydrogels by co-culturing endothelial cells with stromal cells. In order to develop endothelial cell networks as a model of endometrial angiogenesis, HUVECs and HESCs (as stromal cells) were encapsulated into 5 wt% GelMA hydrogels. Encapsulating endothelial (HUVEC) and stromal (HESC) cells in gelatin hydrogels resulted in vessel-like endothelial cell network formation in 3D. The native endometrium contains a vessel length per branch point ranging from approximately 100 - 200 μm over the course of the menstrual cycle [1, 3, 51]. Here, we report endothelial networks formed within the hydrogel that had average branch lengths of approximately 70 μm. While values reported here are similar to physiological levels, future efforts will exploit technologies we have previously reported to incorporate covalently-immobilized VEGF into the GelMA hydrogel to alter the complexity of perivascular networks to study of brain cancer cell invasion [26]. Such techniques, as well as other methods to generate GelMA hydrogels containing spatial-gradients in immobilized biomolecules [52], may provide the ability to further optimize or spatially manipulate average branch length within the range of 100 and 200 μm.

Blastocyst adhesion onto and invasion into the endometrium have never been observed in humans thus, our knowledge of these early stages of implantation come from rare, histological specimens [1]. The establishment of an epithelial monolayer overlaying an endometrial perivascular niche would provide a tool important for studying trophoblast attachment and invasion. A benefit to the GelMA system described in this study is that gelatin natively presents cell binding motifs in its backbone, enabling us to add cells directly to the material without having to incorporate additional cell adhesion peptides [35]. We observed that primary endometrial epithelial cells seeded onto GelMA hydrogels adhered to the hydrogel surface by 6 hours and remained adhered for 3 days. We observed fast (6 hour) attachment of cells to hydrogels which resulted in the formation of cell clusters stacked on top of each other. We expect that monolayer formation is dependent on initial cell seeding density, attachment time, and time in culture. Future in-depth characterization of epithelial monolayers include immunostaining for characteristic markers of epithelial sheets, including E-cadherin and cytokeratin 18, and detailed studies of monolayer functionality, including transepithelial electrical resistance measurements [35, 53, 54].

Models of trophoblast invasion will require an endometrial model that recapitulates the endometrial microenvironment during the window of receptivity. Endometrial stromal cell decidualization, modulated by steroidal sex hormones and cyclic AMP, must occur for human embryo implantation [1, 2]. Decidualized endometrial stromal cells show a characteristic morphological rounding as well as a characteristic increase in prolactin and IGFBP-1 secretion [1, 2]. We demonstrated that the GelMA hydrogel is capable of supporting endometrial stromal cell differentiation for at least 6 days in response to hormonal cues of 1 μM MPA and 0.5 mM 8-Br-cAMP. This decidualization protocol has been used extensively in literature primarily for 2D cultures [30–32, 35], though some success has been reported inducing stromal differentiation with hormonal cues in 3D environments formed from thermally gelled collagen or crosslinked polyethylene glycol macromers [17, 30, 35, 55, 56]. In our study, stromal cells encapsulated in GelMA hydrogels maintained a decidual phenotype for 6 days in culture, as indicated by their epithelial-like morphology as well as increased secretion of characteristic decidual markers prolactin and IGFBP-1. Further characterization and establishment of hormone responsive hydrogel endometrial models would provide critical insights into endometrial function over the course of the entire menstrual cycle.

Monitoring early processes of trophoblast invasion in pregnancy is essential for understanding what drives successful embryo implantation and pregnancy pathologies associated with mutations in implantation. Although trophoblast invasion and endometrial remodeling are critical for the establishment of a successful pregnancy, much is still unknown regarding what drives trophoblast invasion [6, 57]. Our goal was to demonstrate a 3D trophoblast invasion assay developed using a GelMA hydrogel platform. As existing models of invasion in 2D lack the complexity necessary to provide insights into cellular crosstalk and interactions in native 3D environments, 3D models of trophoblast invasion are becoming increasingly a focus for development [15–17, 58, 59]. Here, we seeded 8,000 HTR-8/SVneo cell spheroids into GelMA hydrogels and quantified their invasiveness over the course of 7 days. HTR-8/SVneo cells invaded the GelMA hydrogel matrix over the course of 7 days of culture without exogenous stimulation from various biochemical cues. Studies of this nature provide insight into trophoblast invasion in 3D and allow us to develop invasion assays of increased complexity. We will incorporate promotors (e.g. human chorionic gonadotrophin) and inhibitors (e.g. the transforming growth factor β family) into the cultures and quantify changes in invasion distance in future studies [60–63].

Finally, the efforts reported here used established cell lines, with the exception of the primary endometrial epithelial cells derived from healthy female donors. The use of cell lines for the establishment of this model reduced the cost of the cell sources and provided reproducibility in our studies. Although cell lines provide reproducibility and insight into biological function, primary cells derived from donors would provide the most accurate depiction of endometrial function. HUVECs were used as an endothelial cell source because HUVECs are one of the most widely used cell sources for microphysiological angiogenesis models [64–66]. Interestingly, when using HUVECs in co-culture with endometrial stromal cells, we observed less endothelial network formation compared to studies using HUVECs in co-culture with stromal cells derived from other tissues, such as normal lung fibroblasts [26]. This suggests that cellular crosstalk between HUVECs and HESCs might affect endothelial network complexity. Other labs have demonstrated that although a stromal component featuring stromal cells or other support cells (pericytes, astrocytes, etc.) is necessary for the establishment of stable endothelial networks, specific stromal features, including their tissue source, expression of soluble factors, and mechanical activity, can inhibit or promote vascularization [67–69]. While increasing local presentation of VEGF within the GelMA hydrogel could increase the extent of the endothelial cell network formation, there is also potential to utilize endometrial-specific endothelial cells to create a tissue-specific model of endometrial angiogenesis and alter stromal cell decidualization status to determine the effects of stromal differentiation on endothelial network complexity.

## 5. Conclusions

We report a gelatin hydrogel platform that replicates features of the endometrial microenvironment in GelMA hydrogels. Studies of this nature provide insight into endometrial function and will enable us to study other relevant diseases and processes that occur in the endometrium and during pregnancy. Such tissue engineering methods allow us to rigorously define the degree of biomaterial complexity required to gain functional insight into processes associated with the influence of biomolecular signals, matrix biophysical cues, and heterotypic cell-cell interaction in endometrial physiology and trophoblast invasion. The 3D endometrial platform provides the ability to alter endometrial-relevant features, notably: (i) matrix stiffness, (ii) endometrial angiogenesis, (iii) hormonal stimulation of endometrial stromal cells (e.g. decidualization), (iv) epithelial cell stratification, and (v) trophoblast invasion through a matrix. With these studies, we provide a series of techniques to facilitate future development of endometrial models of increasing complexity. Development of such endometrial models will be transformative for studies pertaining to reproductive health.

## Ethics

This study was not performed with any human subjects or animal subjects.

## Data accessibility

Data are available upon request.

## Author’s contributions

S.G.Z. carried out all lab work, performed all subsequent data analysis, and drafted the manuscript. K.B.H.C. and B.A.C.H. conceived of the study, assisted in experimental planning and analysis, and helped write the manuscript. All authors gave their final approval for publication.

## Funding

Research reported in this publication was supported by the National Institute of Diabetes and Digestive and Kidney Diseases of the National Institutes of Health under Award Numbers R01 DK099528 (B.A.C.H), as well as by the National Institute of Biomedical Imaging and Bioengineering of the National Institutes of Health under Award Numbers R21 EB018481 (B.A.C.H.). The content is solely the responsibility of the authors and does not necessarily represent the official views of the NIH. The authors are also grateful for additional funding provided by the Department of Chemical & Biomolecular Engineering and the Carl R. Woese Institute for Genomic Biology at the University of Illinois at Urbana-Champaign.

### Acknowledgements

The authors would like to acknowledge Ishita Jain (U. Illinois) for her assistance with statistical analysis via R and Dr. Sara Pedron (U. Illinois) for her assistance with NMR analysis.

## Supplemental Materials

**Supplemental 1.** Live/dead images for decidualization experiments.

**Supplemental 2.** Outlier analysis on network metrics.

