## Supplemental Information for "A Gelatin Hydrogel to Study Endometrial Angiogenesis and Trophoblast Invasion"

^1^ Dept. of Bioengineering

^2^ Dept. of Anthropology

^3^ Beckman Institute for Advanced Science & Technology

^4^ Dept. of Chemical and Biomolecular Engineering

^5^ Carl R. Woese Institute for Genomic Biology

University of Illinois at Urbana-Champaign

Urbana, IL 61801

***Supplemental 1. Live/dead images for decidualization experiments.***

After 6 days of exposure to decidualization hormones (1 μM MPA and 0.5 mM 8-Br-cAMP) in cell growth medium (HESC medium) or co-culture medium (EGM), samples (n=2-3; also untreated controls) were stained using calcein, AM (live) and ethidium homodimer 1 (dead) for 20 minutes at 37°C. Samples were then rinsed with PBS then imaged using a Leica DMI 4000 B microscope (Leica Microsystems). Live and dead images were imaged using the same image conditions at the same location. We observed no obvious differences in cell viability between HESC medium and EGM conditions. Treated samples contained very few dead cells.

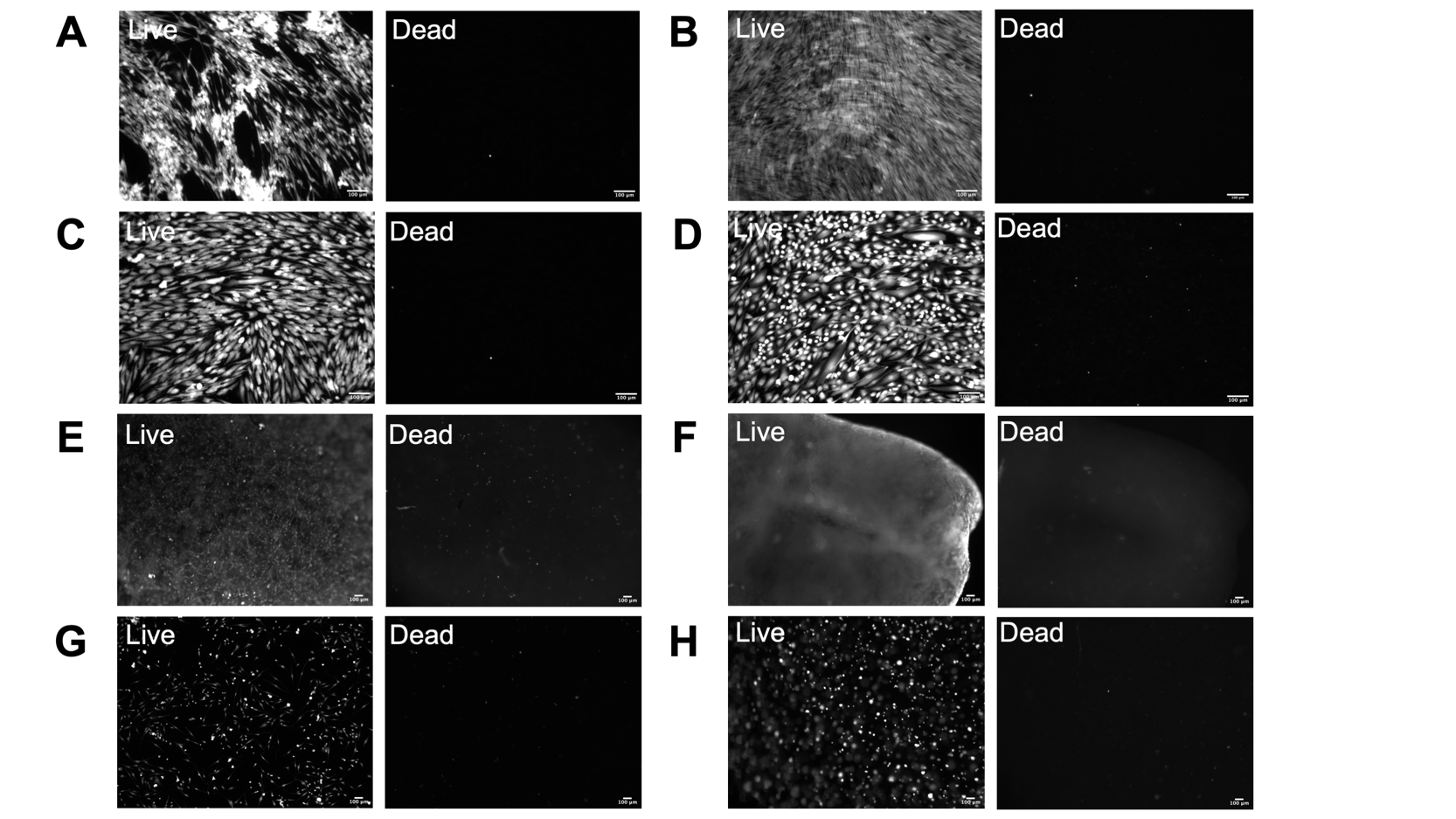

**Supplemental Figure 1.** Representative live and dead images for 2D and 3D control and decidualized HESCs at day 6. HESCs were cultured in growth medium (HESC medium) or co-culture medium (EGM). (*a*-*d*) 2D samples on tissue culture wells: (*a)* Control HESCs cultured in HESC medium. (*b*) Control HESCs cultured in EGM. (*c*) Treated HESCs cultured in HESC medium. (*d*) Treated HESCs cultured in EGM medium. (*e*-*h*) HESCs seeded in hydrogels: (*e*) Control HESCs in a hydrogel cultured in HESC medium. (*f*) Control HESCs in a hydrogel cultured in EGM. (*g*) Treated HESCs in a hydrogel cultured in HESC medium. (*h*) Treated HESCs in a hydrogel cultured in EGM. Scale bars: 100 μm.

***Supplemental 2. Outlier analysis on network metrics.***

Outlier identification and removal was performed in R using rosnerTest from EnvStats v2.3.1 assuming the number of outliers (k) was 3 and α=0.05. The original data set and data set with outliers removed are shown.

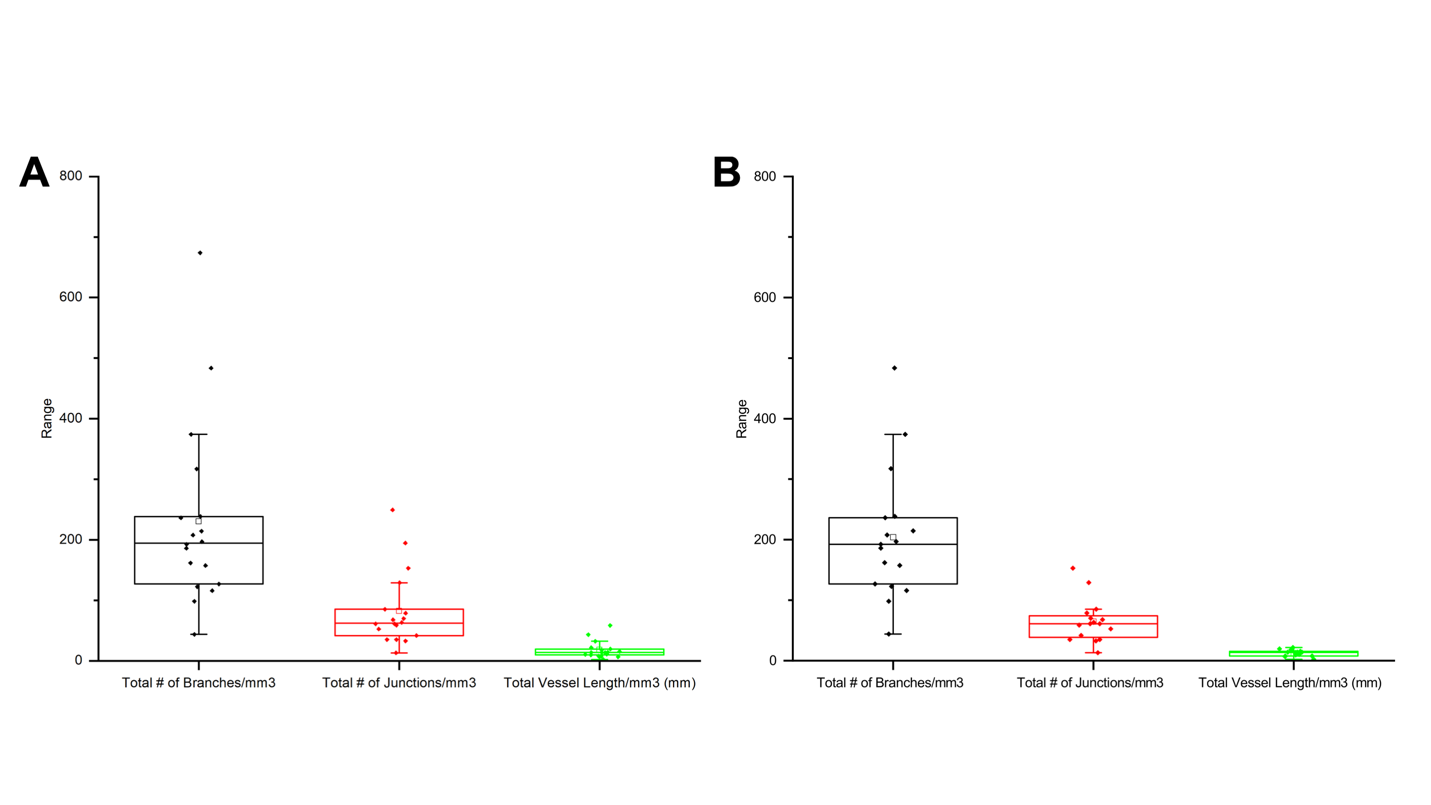

**Supplemental Figure 2.** Boxplots of image analysis metrics for 2:1 HUVEC:HESC ratio samples. (*a*) All data including outliers. (*b*) Data with outliers removed. Outliers were identified and removed using Rosner’s generalized extreme Studentized deviate test assuming 3 suspected outliers and with α set to 0.05. Outliers removed for total # of branches/mm^3^: k=1. Outliers removed for total # of junctions/mm^3^: k=2. Outliers removed for total vessel length/mm^3^: k=3. ☐: mean.
